# Alzheimer’s disease-specific cytokine secretion suppresses neuronal mitochondrial metabolism

**DOI:** 10.1101/2023.04.07.536014

**Authors:** Madison K. Kuhn, Rebecca M. Fleeman, Lynne M. Beidler, Amanda M. Snyder, Dennis C. Chan, Elizabeth A. Proctor

**Affiliations:** Department of Neurosurgery, Penn State College of Medicine, Hershey, PA, USA; Department of Pharmacology, Penn State College of Medicine, Hershey, PA, USA; Department of Biomedical Engineering, Pennsylvania State University, University Park, PA, USA; Department of Engineering Science & Mechanics, Pennsylvania State University, University Park, PA, USA; Center for Neural Engineering, Pennsylvania State University, University Park, PA, USA; Department of Microbiology & Immunology, Penn State College of Medicine, Hershey, PA, USA; Department of Neurology, Penn State College of Medicine, Hershey, PA, USA

**Keywords:** neuroinflammation, immunometabolism, multivariate modeling, ATP production, mitochondrial respiration

## Abstract

**Introduction:** Neuroinflammation and metabolic dysfunction are early alterations in Alzheimer’s disease brain that are thought to contribute to disease onset and progression. Glial activation due to protein deposition results in cytokine secretion and shifts in brain metabolism, which have been observed in Alzheimer’s disease patients. However, the mechanism by which this immunometabolic feedback loop can injure neurons and cause neurodegeneration remains unclear.

**Methods:** We used Luminex XMAP technology to quantify hippocampal cytokine concentrations in the 5xFAD mouse model of Alzheimer’s disease at milestone timepoints in disease development. We used partial least squares regression to build cytokine signatures predictive of disease progression, as compared to healthy aging in wild-type littermates. We applied the disease-defining cytokine signature to wild-type primary neuron cultures and measured downstream changes in gene expression using the NanoString nCounter system and mitochondrial function using the Seahorse Extracellular Flux live-cell analyzer.

**Results:** We identified a pattern of up-regulated IFNγ, IP-10, and IL-9 as predictive of advanced disease. When healthy neurons were exposed to these cytokines in proportions found in diseased brain, gene expression of mitochondrial electron transport chain complexes, including ATP synthase, was suppressed. In live cells, basal and maximal mitochondrial respiration were impaired following cytokine stimulation.

**Conclusions:** An Alzheimer’s disease-specific pattern of cytokine secretion reduces expression of mitochondrial electron transport complexes and impairs mitochondrial respiration in healthy neurons. We establish a mechanistic link between disease-specific immune cues and impaired neuronal metabolism, potentially causing neuronal vulnerability and susceptibility to degeneration in Alzheimer’s disease.

## Introduction

Alzheimer’s disease (AD) is a progressive neurodegenerative disorder resulting in cognitive decline and memory loss. AD is estimated to affect more than 6.5 million people in the United State and 55 million worldwide, with prevalence predicted to increase with the world’s growing aged population.^108^ There are no proven therapies to prevent or cure the disease, which is in part due to an incomplete understanding of the disease’s complex etiology. AD is histologically characterized by neuron and synapse loss, reactive gliosis, and misfolded aggregated proteins of extracellular amyloid beta (Aβ) plaques and intracellular neurofibrillary tangles.^78^ Neuroinflammation is an early event in AD,^15,40^ appearing before the development of clinical symptoms^13,83^ and often prior to the detection of proteinopathy in AD mouse models.^39,91,101^ Neuroinflammation is an active driver of disease and is sufficient to initiate neurodegeneration.^40^ Importantly, the majority of known AD risk genes participate in immune response.^38,68^ Immunomodulatory therapies are gaining interest for their potential to interfere with disease progression or even disrupt disease initiation. However, the entangled nature of the immune response complicates the identification of robust, effective targets that will not detrimentally affect immune signals necessary for brain health.^40^ In this study, we examine the evolution of cytokine expression over aging and disease progression in a transgenic mouse model of AD (5xFAD)^62^ and wild-type littermates to generate a systems-level profile of dysregulated immune signaling in the hippocampus, a primary brain region involved in AD onset and progression.^70^ This systems biology approach allows us to identify key AD-implicated cytokine cues and their downstream detrimental effect on neuron health.

Neuronal support cells in the brain, called glia, provide immune regulation that is necessary to maintain brain homoeostasis. However, their prolonged activation can cause damaging and even neurotoxic inflammation.^12,34^ Microglia, the resident immune-competent cells of the brain, play essential roles in development and disease and are thought to be the primary contributor to neuroinflammation in AD.^16,40,95^ Another glial cell type, astrocytes, offer a variety of support functions to neurons including metabolic support and neurotransmitter regulation. Additionally, astrocytes play a role in the immune response, helping to clear debris and amplify microglial signaling cascades.^24,35^ In neurodegenerative disease, beneficial immune responses, such microglial activation for the clearance of debris,^95^ can become detrimental: prolonged microglial activation is accompanied by an overproduction of pro-inflammatory factors, aberrant synapse engulfment, and the release of nitric oxide.^12,37^ Concurrently, dampened immune responses result in ineffective neuroprotection.^52,74^ The shift between detrimental and beneficial roles of immune activation, in combination with the substantial overlap in these roles, complicates the identification of therapeutic targets to ameliorate harmful neuroinflammation.^24,37^ Consequently, these therapies may also have a limited therapeutic window.^37^ Immunosuppressant drugs have not thus far been found to consistently reduce one’s risk for AD or have protective effects in its progression.^39^ However, the potential remains that if upstream, broad neuroinflammatory pathways cannot be modulated with enough specificity to benefit disease state, targeting of downstream pathways contributing to neuronal injury may prove to be more advantageous.

Metabolism deficits in AD are similarly characteristic of early disease and are closely intertwined with the immune response, contributing to pathology progression and neurodegeneration through an immunometabolic feedback loop.^87^ Evidence suggests that metabolic dysregulation precedes proteinopathy and participates in the initiation of AD.^18,81^ The complimentary and synergistic action of neuroinflammation and metabolic impairment in early disease creates a unique opportunity to maximize therapeutic effect by disrupting this immunometabolic feedback interaction – a potentially more efficacious target than immune signaling or metabolic function alone. However, the mechanism by which immunometabolic feedback injures neurons and promotes neurodegeneration remains unclear. Here, we investigate downstream dysregulation of neuronal metabolic pathways as a result of a disease-relevant signature of immune cues to identify specific and actionable neuronal vulnerabilities created by immunometabolic feedback in the hippocampus.

Cytokines, immune signaling molecules, both initiate and are a direct consequence of a variety of immune responses and cell-to-cell immune signaling^6,104^ and are often dysregulated in the AD brain.^74^ Cytokine signaling is highly interdependent, forming a complex web of communications between multiple cell types, and often results in a signaling cascade that prompts the secretion of additional cytokine cues.^104^ These characteristics complicate the untangling of shifting beneficial and detrimental roles of immune responses in neurodegeneration, but highlight the necessity of a holistic view of cytokine signaling to describe the overall immune state of the brain. To identify key pathways implicated in disease, we profiled a broad panel of cytokines and used a multivariate mathematical modeling tool, partial least squares (PLS), to construct a signature of cytokine cues associated with AD progression. The actions and downstream effects of individual cytokines are often measured and investigated independently, leading to an incomplete and potentially incorrect understanding of their actions in concert. We applied our signature of AD-upregulated cytokines to healthy neuron and astrocyte cultures and analyzed the resulting differential gene expression and subsequent activation of pathways affecting neuronal viability and cellular metabolism. We determined that the AD cytokine signature suppressed gene expression of numerous mitochondrial electron transport chain (ETC) complex subunits and impaired the mitochondrial function of living neurons. Exposure to the cytokine signature representing the milieu of the AD brain thus predisposes neurons to injury from inadequate energy metabolism and reactive oxygen species (ROS) damage, which would in turn exacerbate neuroinflammation. Our established mechanistic link between the disease-specific cytokine signaling and impaired neuronal metabolism identifies opportunities to target specific detrimental downstream metabolic pathways to disrupt the complex, cooperative actions of neuroinflammation and dysregulated metabolism in disease.

## Methods

### Mice

All animal procedures were approved by the Penn State College of Medicine Institutional Animal Care and Use Committee (PROTO201800449). 5xFAD (B6SJL, Jackson Laboratory) breeding pairs were purchased for the generation of animals used in the longitudinal study of cytokine profiles over Alzheimer’s disease pathology progression. The 5xFAD strain is hemizygous for 5 familial Alzheimer’s disease mutations under the Thy1 promoter for expression in the brain: the Swedish (K670N, M671L), Florida (I716V), and London (V717I) mutations in human amyloid beta precursor protein (APP) and the M146L and L286V mutations in human presenilin 1 (PSEN1).^62^ Transgenic and noncarrier littermates were housed together and genotyped according to Jackson Laboratory’s standard PCR assay protocol 31769 using DNA isolated from tail clippings. Even numbers of male and female mice (n=5 per sex, per group) were sacrificed for the timepoints of this study. CD1 breeder pairs and untimed pregnant females were purchased from Charles River for the generation of primary cell cultures. All animals were provided with food and water *ad libitum* and maintained under a 12-hour light/12-hour dark cycle.

### Tissue collection and preparation

We aged the mice to 30, 60, 120, and 180 days to profile cytokine concentrations prior to the deposition of intraneuronal Aβ at 1.5 months, at the development of extracellular amyloid and gliosis at 2 months, at the first appearance of synaptic loss and cognitive deficits at 4 months, and with neuronal loss at 6 months in the 5xFAD model.^62^ At each timepoint, 5xFAD mice and noncarrier littermates were sacrificed by unanesthetized decapitation using a mouse guillotine (Braintree Scientific). The brain was isolated and placed into cold HEPES-buffered Hanks’ Balanced Salt solution (pH 7.8). The hippocampus of the right hemisphere was dissected, meninges removed, flash-frozen in liquid nitrogen, and stored at -80°C. After all study samples were collected, hippocampi were thawed on ice and mechanically digested by trituration in 200 μL of RIPA buffer (Boston BioProducts P8340) with protease inhibitor cocktail (Thermo A32953, 1 Pierce mini Protease Inhibitor tablet/ 10 mL RIPA). Digested samples were vortexed vigorously for 1-2 minutes, rested for at least 20 minutes, and then centrifuged at 5000xg for 5 minutes. Supernatants were transferred to new sample tubes for analysis. An aliquot of each homogenate sample was set aside for total protein quantification, and the rest was flash frozen in liquid nitrogen and stored at -80°C for Luminex multiplex immunoassays.

### Quantification of total protein

The total protein content of the hippocampal homogenate samples was quantified using the Pierce BCA Protein Assay Kit (Fisher 23225) according to the manufacturer’s instructions. Samples were run in duplicate and absorbances read using a SpectraMax i3 minimax 300 imaging cytometer (Molecular Devices). Sample concentrations were quantified by linear regression using duplicate standard samples ran on each plate.

### Luminex multiplex assays

Cytokine concentration levels of 5xFAD hippocampal homogenate samples were quantified using the Milliplex Map Mouse Cytokine/ Chemokine Magnetic Kit (Millipore MCYTMAG-70K-PX32), measuring a broad panel of secreted cytokines, on a the Luminex FLEXMAP3D platform. The assay was performed according to the manufacturer’s protocol with minor modifications to accommodate the use of 384-well plates. The magnetic beads and antibody solutions were diluted 1:1 and used at half volume, and the streptavidin-phycoerythrin was used at half volume. Equal amounts of RIPA buffer and assay buffer were used in the preparation of the standards and blanks. Homogenate samples were thawed and stored on ice. Samples were diluted to 0.3 g/mL total protein using RIPA buffer, allowing 7.5 μg total protein per well, and assayed in technical triplicate.

### Cytokine profile data cleaning

The Luminex Xponent software was used to interpolate sample cytokine concentrations from 5-point logistic standard curves. Concentrations below the detection limit (< 3.2pg/mL) were assigned 0 pg/mL. The raw concentration data was processed using an automated in-house pipeline, available from GitHub at https://github.com/elizabethproctor/Luminex-Data-Cleaning (Version 1.02). Briefly, the pipeline removes readings generated from less than 20 beads (4 observations in the current dataset) and calculates the pairwise differences of the remaining technical triplicates. If the difference between one replicate is greater than twice the distance between the other two, that replicate is removed from the analysis. The average of the remaining technical replicates for each cytokine is then calculated for the final dataset for use in subsequent multivariate analyses. Following the automated data cleaning, we removed entire cytokines from the dataset if over half the readings were 0 pg/mL in a pattern that did not partition between experimental groups, such as disease or timepoint.

### Partial least squares modeling

Cytokine signatures were constructed using the linear, supervised multivariate mathematical modeling tool partial least squares (PLS).^33,94^ We chose PLS to account for the highly interdependent nature of cytokine expression and signaling. PLS allows for the identification of significant multivariate changes in correlative predictors (here, the cytokines) as they relate to a dependent response or group (e.g., timepoint). This aim is achieved by building linear combinations of the predictors (latent variables, LVs) that maximize the covariation between the predictors and the response. This maximization of multivariate covariance in predictors with response allows us to identify subtle but meaningful patterns in cytokine expression that are undetectable using univariate analysis methods that are unsuitable for highly correlative variables.

Partial least squares regression (PLSR) was conducted for prediction of continuous numerical outcomes, such as timepoint, while partial least squares discriminant analysis (PLS-DA) was used in the discrimination of experimental groups, such as disease versus control. PLS models were built in R using the *ropls* package^84^. Cytokine data were mean-centered and unit-variance scaled prior to PLS modeling. To generate each model, the optimal number of latent variables was determined by repeated random sub-sampling cross-validation. Cross-validation test sets were randomly generated using 1/3 of the dataset if the number of samples used to build the model was >30 or 1/5 of the dataset if samples <30. Cross-validation was repeated 100 times. For every iteration, models were built using the training set (data excluded from test set) comprised of either 1, 2, 3, 4, or 5 LVs. The resulting models were used to predict the group identity or response value of each sample in the test set to estimate classification accuracy (in PLS-DA) or root mean squared error of cross validation (RMSECV) (in PLSR). RMSECV was calculated by:

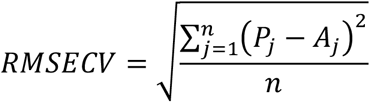

Where n is the number of samples in the test set, P is the predicted value, and A is the actual value of the jth sample. The test/training set was randomly regenerated for the following iterations, and the average of the estimated classification accuracy or RMSECV following cross-validation was calculated for each of the 1, 2, 3, 4, or 5 LV PLS models. The model with the number of LVs resulting in the best accuracy/ lowest error was chosen for subsequent analysis and significance testing. A model’s significance was calculated using a permutation test. For each iteration of the permutation test, the group identities or responses were randomly shuffled among samples, preserving the data landscape of the cytokine profiles while creating randomized associations. Cross-validation was performed as described previously, preserving the number of latent variables comprising the “real” model, to estimate the true random accuracy. The p-value of the optimized experimental model was then calculated by comparison with the mean and standard deviation of the distribution of random models’ accuracy. For PLS models containing 1 latent variable, the second latent variable is shown in the scores plot for ease of visualization but was not included in the calculation of error, p-value, or Variable Importance in Projection (VIP) scores. Models presented in this study were orthogonalized for improved interpretability. Orthogonalization maximally projects covariation of the measured cytokines with the response to the first latent variable, prioritizing this covariation over variation in the measured cytokines between samples.^86^ We identified key differential cytokines by their variable importance in projection (VIP) score, a measure of a variable’s normalized contribution to the predictive accuracy of the model across all latent variables.^32^ Loadings on the first latent variable of cytokines with VIP score > 1, indicating greater than average contribution to the model, are designated as the cytokine signature of the given outcome (e.g. AD progression).

### Primary cell culture

Primary hippocampal neuron and astrocyte cultures were generated from P0 CD1 neonates. Pups were sacrificed by decapitation using surgical scissors, and the brains were removed and placed in pre-chilled sterile dissection medium (HEPES buffered Hanks’ Balanced Salt solution, pH 7.8) on ice. Hippocampi were isolated and placed into a sterile Eppendorf tube containing cold dissection medium following the removal of meninges. For each cell culture experiment, hippocampi from 6-8 neonates were combined. The tube was centrifuged briefly, and the dissection medium was aspirated. Warm neuronal plating medium (Neurobasal Plus, 10% FBS, 1X GlutaMAX, 1X Penicillin-Streptomycin (Gibco): for neuron cultures) or glial medium (DMEM/F12, 20% FBS, 1X Penicillin-Streptomycin, 1mM Sodium Pyruvate (Gibco), 10 ng/mL Epithelial Growth Factor: for astrocyte cultures) was added to the tube. The sample was manually triturated with a p1000 pipet followed by a p200 pipet.

For neuron cultures, the total cell concentration was measured using the Countess II automated cell counter (Invitrogen), and 5.47*10^5^ cells/cm^2^ were plated in each well of a poly-D-lysine-coated Seahorse XFe24 V7 PS Cell Culture Microplate (Agilent) or a 24-well tissue culture plate. Following cell attachment 2-4 hours after neuron plating, plating medium was removed and neuronal culture medium was added (Neurobasal Plus, 1X B27 Plus supplement, 1X GlutaMAX, 1X Penicillin-Streptomycin (Gibco)). Half of the neuronal culture medium was changed after 5 days, and all the neuronal medium was changed on day 9-10 with the addition of experimental treatments. Neuron cultures were assayed or lysed on day 12-13.

For astrocyte cultures, the cell solution was transferred to a poly-D-lysine (Gibco)-coated tissue culture flask with additional glial medium. Glial medium in astrocyte culture flasks was replaced every 2-3 days until cells reached confluency. Flasks were then placed on an orbital shaker in a cell culture incubator for 2 hours at 180 rpm, and the cell culture medium containing non-adherent cells was removed. The remaining adherent astrocytes were washed with 1X PBS, detached using 0.05% trypsin-EDTA, quenched with fresh glial medium, and evenly distributed into the wells of a poly-D-lysine-coated 6-well tissue culture plate. Glial medium was switched every 2-3 days and treatments were applied at 50% confluency in the 6-well plates. All cell culture plates (neurons and astrocytes) were maintained at 37°C and 5% CO_2_.

### Cytokine stimulation of primary culture

The up-regulated Alzheimer’s disease-specific cytokine profile of IFNγ, IP-10, and IL-9, as determined by the multivariate modeling, was applied to primary neuron and astrocyte cultures derived from CD1 P0 neonates. The combination cytokine treatment was applied 72 hours prior to experimentation in levels proportional to the concentrations we measured in 5xFAD 180-day hippocampus samples. Concentrations were centered in the nanomolar range previously used to study acute cytokine responses in neuron cultures.^23,89^ Recombinant murine cytokines were purchased from Peprotech, IFN-γ (cat 315-05), IP-10 (cat 250-16), and IL-9 (cat 219-19), and reconstituted to 1 mg/mL in sterile water and diluted to 250 μg/mL (IL-9) or 10 μg/mL (IFNγ and IP-10) in 0.1% cell-culture grade bovine serum albumin (Sigma A9418) in 1X PBS. 5nM IFNγ, 12nM IP-10, and 500nM IL-9 or an equal amount of 0.1% bovine serum albumin vehicle were added to the appropriate neuronal or glial cell culture medium for treatment. Neuron cultures were treated 9 or 10 days after plating, and astrocytes were treated, following the removal of non-adherent cells, at 50% confluency. After a 72-hour stimulation, cells were assayed on the Seahorse XFe24 Extracellular Flux Analyzer or lysed for RNA isolation.

### RNA isolation and NanoString nCounter analysis

Cytokine- or vehicle-stimulated primary CD1 astrocyte cultures in 6-well plates and neuron cultures in 24-well plates were lysed and their total RNA extracted using RNeasy Mini Kits (Qiagen 74104) according to the manufacturer’s instructions. The lysis of 2 wells of astrocytes or 3 wells of neurons were combined for each biological replicate to minimize technical variation, resulting in N=3 biological replicates for each treatment/cell-type. The RNA content and quality of each sample was determined using the NanoDrop 2000 Spectrophotometer (Thermo). RNA samples had a 260/280 absorbance ratio of 1.9 or greater and a 260/230 ratio of 1.8 or greater, quality control guidelines outlined by NanoString to exclude samples containing protein or organic compound contamination. Targeted gene expression was quantified using NanoString nCounter technology and the Mouse Metabolic Pathways Panel, with the assay performed by the NanoString Proof of Principle Lab (Seattle, Washington) to measure expression of 768 metabolism-relevant genes and 20 common housekeeping genes. The raw expression data were processed using the NanoString nSolver Analysis software. All samples passed quality control (QC) with the default parameters. Data normalization was performed separately for neuron and astrocyte samples. For background thresholding, genes with a raw count less than 20 were reassigned a threshold count of 20, in order to calculate fold changes to the same baseline reading. Data was normalized using a 2-step positive control and housekeeping gene normalization. For the selection of appropriate housekeeping genes for each cell type, housekeeping genes with counts under 100 were first removed. Then, the housekeeping genes with a coefficient of variation of 10% or less were kept for normalization (5 genes kept for neurons, 6 genes for astrocytes). Using the final normalized astrocyte and neuron expression data, the log2 fold changes of cytokine versus vehicle treatment comparisons were calculated using the nSolver software. Genes with a significant log2 fold change, as determined by a p-value of 0.05 or less, were kept for further analysis. Significantly differentially expressed genes were excluded if the average counts for both cytokine and vehicle treatments were both below 30, close to the assigned background threshold of 20.

### Construction of protein-protein interaction networks

Significantly differentially expressed genes in cytokine-treated neurons and astrocytes were used to construct protein-protein interaction networks using the STRING database.^82^ Pair-wise gene inputs were given a score based on known and predicted protein interactions from experimental, co-expression, and gene fusion evidence. The Cytoscape StringApp was used for network visualization^22^ with the edge-weighted, spring-embedded layout. The nodes of the networks are individual significantly differentially expressed genes. The genes were characterized by their overall functional group.^57^ Isolated sub-networks of 4 or fewer genes were removed.

### Seahorse Mito Stress Test

Neuronal mitochondrial function following vehicle or cytokine treatment was characterized on the Seahorse XFe24 Extracellular Flux Analyzer platform (Agilent). The Seahorse Analyzer measures oxygen consumption rate (OCR) to quantify characteristics of mitochondrial function. During the Mito Stress test, a series of compounds is injected into the cell culture medium to inhibit complexes of the mitochondrial electron transport chain (ETC) to isolate readings of basal respiration, maximal respiration, ATP production, and non-mitochondrial respiration. Following baseline oxygen consumption readings (basal respiration), oligomycin A, an inhibitor of ATP synthase, is added the cell culture medium and the subsequent drop in oxygen consumption corresponds to the mitochondrial ATP production by ATP synthase. The following addition of FCCP, an uncoupling agent that disrupts the mitochondrial membrane potential, allows for uninhibited electron flow, and the resulting increase in oxygen consumption demonstrates the maximal respiration of the mitochondria. Finally, rotenone and antimycin A (inhibitors of complex I and III, respectively, and together considered a mitochondrial poison) are added to disable the electron transport chain and, thus, residual consumption of oxygen is driven by non-mitochondrial processes.

Primary CD1 hippocampal neurons were stimulated with 5 nM IFNγ, 12 nM IP-10, and 500 nM IL-9 for 72 hours prior to Seahorse assays. Neuron culture wells were washed with Seahorse assay medium and given fresh assay medium: DMEM (cat 103680-100), 10 mM glucose solution (cat 103577-100), 1 mM pyruvate solution (cat 103578-100), 2 mM glutamine solution (cat 103579-100) (Agilent). Neuron cultures were transferred to a non-CO_2_ incubator for 1 hour prior to running the assay. All Seahorse compound stocks were diluted to 50 mM working stocks with assay medium and added to the appropriate Seahorse cartridge ports to give final well concentrations of 4 μM oligomycin A, 4 μM FCCP, and 1 μM of both rotenone and antimycin A after each injection. Following the assay, the neuron culture wells were washed with 1X PBS, and the cells were lysed in 25 μL lysis buffer (Millipore 43-040) by manual titration. The total protein content of each lysis sample was quantified using the Pierce BCA Protein Assay Kit (Fisher 23225) according to the manufacturer’s instructions in technical triplicate with standards measured on each plate. Total protein content was used to normalize intra-plate variation using the Seahorse Wave software. Individual wells were excluded from subsequent analysis if the cells were not viable (i.e. near-zero OCR measurements) or did not appropriately respond to the compounds (e.g. FCCP did not increase oxygen consumption). Each biological replicate was an independent Seahorse experiment of neurons derived from a unique group of neonates and was the average measurement (calculated according to Table 1) of the readings of 5-10 wells. Final OCR measurements of the biological replicates were normalized to non-mitochondrial respiration to account for batch effects.^60^ Two-tailed Student’s T Tests were conducted to determine significant differences in basal respiration, maximal respiration, ATP production, and proton leak in OCR measurements between cytokine and vehicle treatments. Statistical significance was determined by a significance level of less than 0.05.

**Table 1:** Calculations of Seahorse Mito Stress Test Assay Measurements.

| Measurement | Calculation |
| --- | --- |
| Non-mitochondrial respiration | Median OCR measurement following rotenone/antimycin A injection |
| Basal Respiration | (Median OCR measurement prior to the addition of oligomycin A) – (Non-mitochondrial respiration) |
| ATP production | (Basal respiration) – (minimum OCR measurement following oligomycin A injection) |
| Maximal Respiration | (Maximum OCR measurement following FCCP injection) – (Non-mitochondrial respiration) |
| Proton Leak | (Minimum OCR measurement following oligomycin A injection) – (Non-mitochondrial respiration) |

## Results

### The cytokine signature of AD is distinct from that of healthy aging

To define the evolution of the immune milieu of the hippocampus over the course of disease progression in the 5xFAD model, we first built a partial least squares regression (PLSR) model to regress timepoint against 5xFAD hippocampal cytokine concentrations (Fig. 1A). We identified the above-average contributors to the predictive accuracy of our model (2 latent variables, RMSECV=58.4, p=10^−5^) using the variable importance in projection score (Methods). We found that IP-10, IL-9, and IFN-γ, are up-regulated over the course of disease, and levels of IL-2 and IL-1α decrease with disease progression. This signature describes a milieu that enacts a two-pronged increase in neuronal vulnerability: first, by increasing expression of canonically pro-inflammatory cytokines, ^4,45,55,56,79,80,88^ and second, by decreasing expression of cytokines with neuroprotective functions in the brain. ^72,73,102^ However, subtleties exist in these relationships, some cytokines having both pro- and anti-inflammatory actions, and others whose action is incompletely understood in the brain or in the context of AD.^14,17,23,56,59,93^

**Fig 1.**
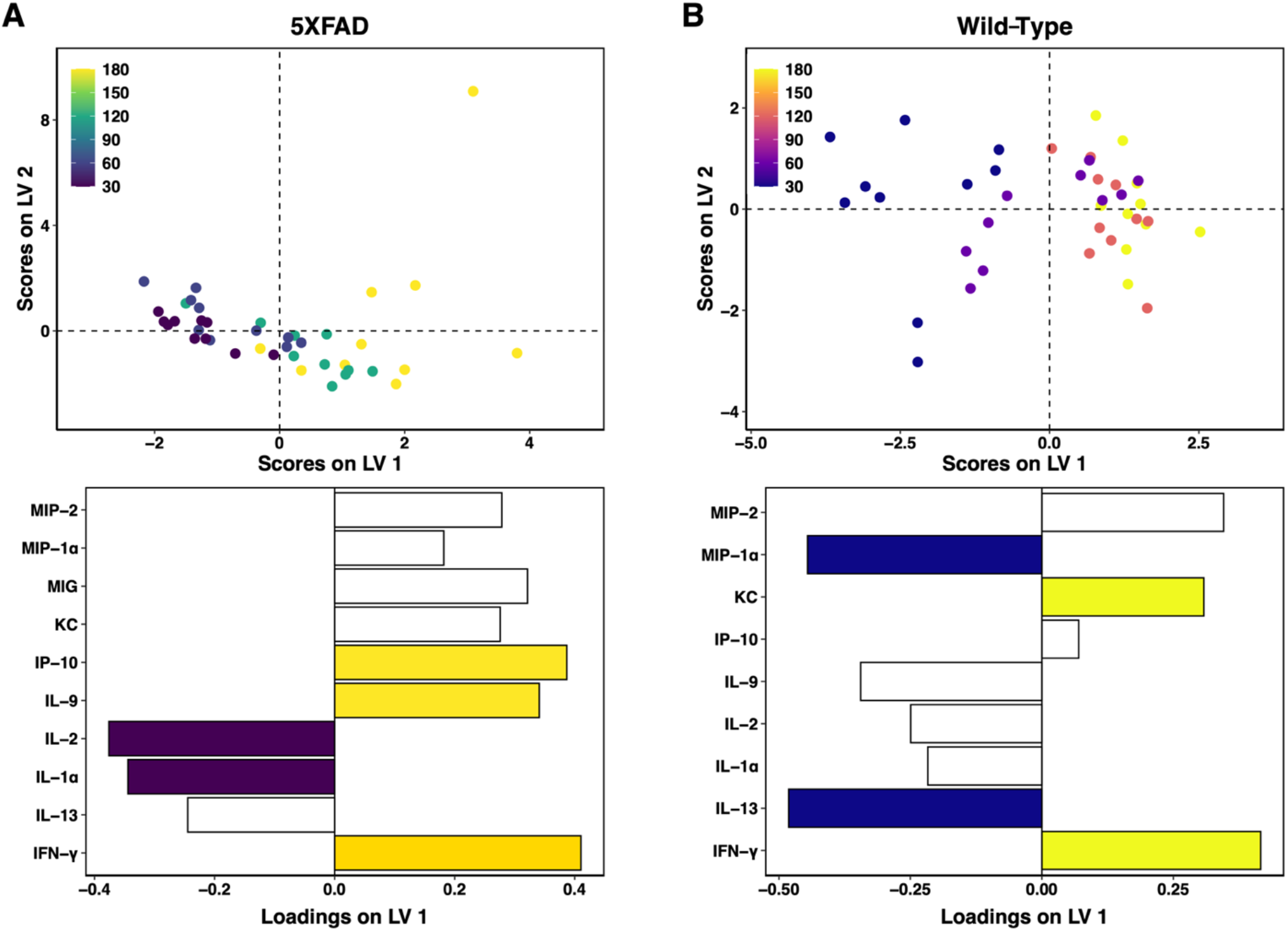
Unique cytokine signatures predict aging in 5xFAD and wild-type hippocampi. Scores plot (top) and LV1 loadings plot (bottom) of PLSR models regressing cytokine concentrations against timepoint in (A) 5xFAD hippocampus (2 LV, RMSECV: 58.4, p-value: <0.00001) and (B) wild-type hippocampus (1 LV, RMSECV: 51.9, p-value: 0.00001). Colored loadings signify cytokines with a VIP score > 1. Each point represents a single mouse. PLS modeling is comparative and not absolute: older mice (positive scores on LV1) feature up-regulation of cytokines with positive loadings and down-regulation of cytokines with negative loadings, as compared to younger mice (negative scores on LV1).

To distinguish the evolving milieu of immune cues in AD onset and progression from that of normal aging, which also exhibits an immune component (“inflammaging”),^10,28,29^ we constructed a PLSR model regressing timepoint against cytokines profiles of wild-type samples (Fig. 1B). The key contributors to the predictive accuracy of the model (1 latent variable, RMSECV=51.9, p=10^−5^) are increased IFN-γ and KC/CXCL1 and decreased MIP-1α and IL-13 with advancing age. Notably, the up-regulated immune cues in healthy aging and AD are largely non-overlapping, indicating that AD onset and progression involves pathological changes in a distinct set of process from those that change in the normal course of aging. Emphasizing these distinct processes, MIG/CXCL9, while not a key contributor to our AD signature, was identified as increasing with disease progression, but was absent in wild-type mice at all timepoints. In contrast, because the majority of cytokine species were down-regulated with aging in the wild-type model, all down-regulated species in the AD signature discussed above are also decreasing in healthy aging, suggesting a natural decrease in immune responsiveness or surveillance. However, we note that a key decreasing cytokine in healthy aging, MIP-1α/CCL3, increases with disease, although not as a key contributor to the AD signature. We conclude that while similarities exist between the immune milieus of healthy aging and AD, which is fitting for a disease with the primary risk factor of age, they are minimally overlapping, supporting a fundamental shift in the brain rather than accelerated aging, as has been suggested by some.^99^

These models independently assess how cytokine levels change with the progression of aging or of AD. While there are distinct differences in the disease and wild-type models, particularly in the contributions of IP-10, IL-9, and MIP-1α, there is also overlap, and any comparison of such separately-constructed models is necessarily qualitative. We observed cytokines implicated in AD, such as CXCL1 and IL-13,^30,54,66,98,105^ to also be important for the prediction of age in wild-type samples. In order to directly and quantitatively compare the hippocampal immune milieu in AD or normal aging, we next constructed PLS models to differentiate the AD and wild-type groups at each.

### AD mice exhibit neuroimmune over-activation late in the disease timeline

To directly compare cytokine signatures of AD and wild-type control mice, we constructed partial least square discriminant analysis (PLS-DA) models to predict disease status at 30 or 180 days, representing prodromal and fully symptomatic AD, respectively. In 30-day-old hippocampus, prior to intracellular deposition of Aβ, we discriminate AD from wild-type with 83% accuracy and high significance (p-value: <0.005, Fig. 2A). The strongest contributors to predictive accuracy, MIP-1α and IL-9, are both down-regulated in AD before the development of pathology, in comparison to healthy controls. When we compare this result with the AD- and aging-specific progression signatures (Fig. 1), we find that both MIP-1α and IL-9 decrease with healthy aging but increase with disease progression.

**Fig 2.**
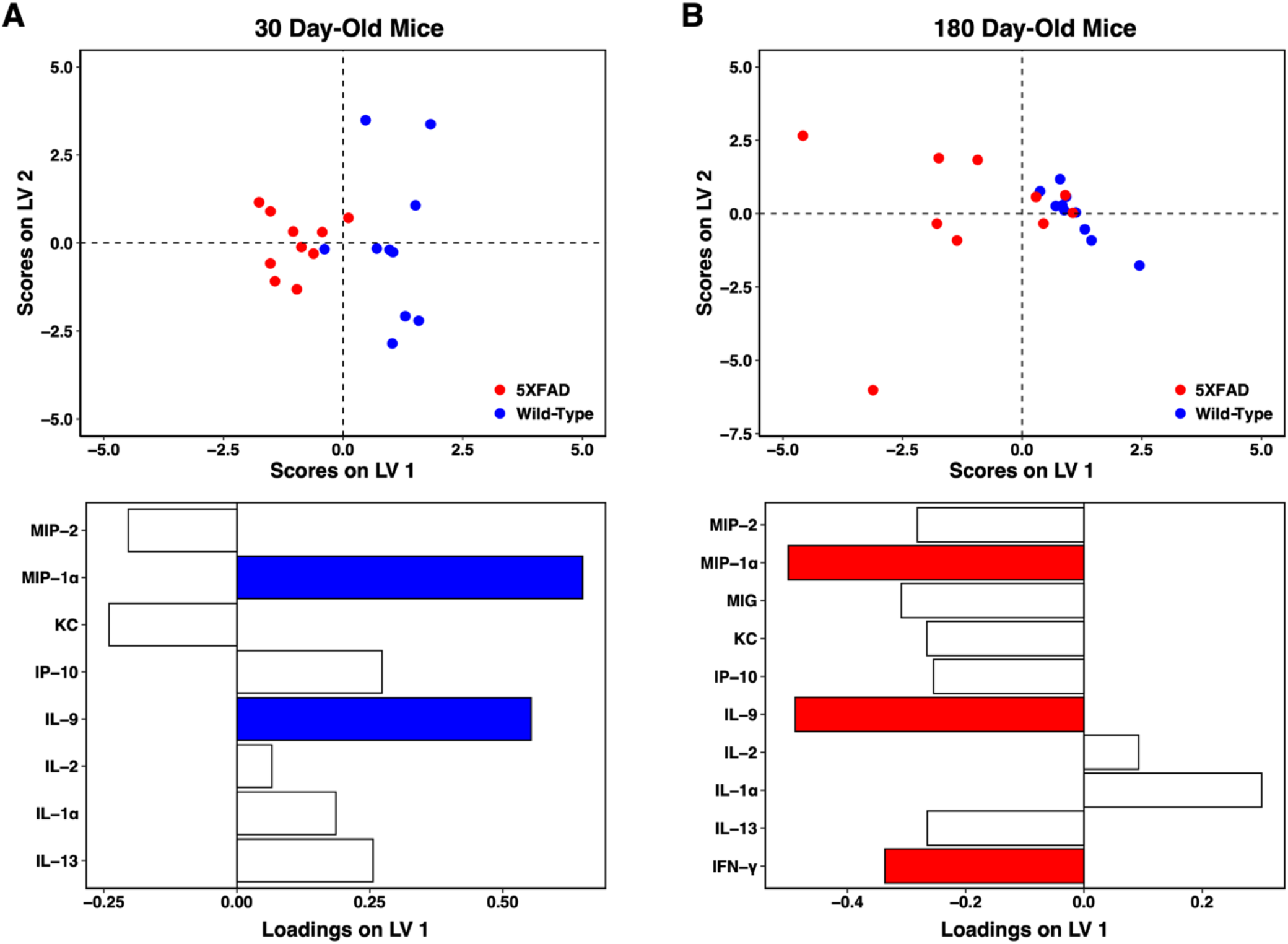
Cytokines are up-regulated in aged 5xFAD hippocampus but dampened in comparison to wild-type prior to protein deposition. PLS-DA scores plot (top) and LV1 loadings plot (bottom) of (A) 30-day 5xFAD vs wild-type hippocampus (3 LV, accuracy: 83%, p-value: <0.005) and (B) 180-day 5xFAD vs wild-type hippocampus (2 LV, accuracy: 80%, p-value: <0.01). Colored cytokine loadings signify cytokines with a VIP score > 1. Each point represents a single mouse. PLS modeling is comparative and not absolute: 5xFAD mice (negative scores) feature down-regulation of cytokines with positive loadings and up-regulation of cytokines with negative scores, as compared to wild-type mice (positive scores on LV1).

The 180-day PLS-DA model features a similarly high predictive accuracy (80%) and significance (p-value: <0.01) in discriminating cytokine expression in AD versus wild-type control hippocampus. The model describes an overall increase in cytokine levels in late-stage AD as compared to healthy controls. The strongest contributors to predictive accuracy were IL-9, IFN-γ, and MIP-1α, all up-regulated in AD. These results are in agreement with the AD- and aging-specific progression signatures, where IL-9 and MIP-1α both increased with AD progression but decreased with healthy aging. While IFN-γ increased in both AD progression and healthy aging, this direct comparison of AD with wild-type control at the 180-day time point demonstrates that the increase in disease is significantly more pronounced than that in healthy aging, a point that cannot be drawn from the comparison of the two separately-derived models in Figure 1. A similar point can be drawn for KC/CXCL1, which was also up-regulated in both AD progression and healthy aging but is significantly higher in disease samples at 180 days.

In total, these models reflect low immune activity at an early age in the AD brain, which may suggest an increased vulnerability to disease. With the onset of disease pathology and fully symptomatic AD, the AD brain exhibits a significant increase in overall cytokine expression as compared to wild-type control, with a specific pro-inflammatory signature that agrees with the time-evolution signature of AD progression when considered outside of comparison to healthy aging.

### AD cytokine signature alters metabolic gene expression in neurons but not astrocytes

The cytokine signature of AD progression represents the evolving neuroimmune milieu in the AD brain. We next endeavored to characterize the effect of this evolving milieu on neuron health. We approached this question from two directions: the direct effect of the AD cytokine signature on neurons, and the indirect effect of loss of support to neurons *via* astrocyte dysfunction.^7,48^ Astrocytes provide critical metabolic support to neurons, and disruption of this support would lead to increased neuronal vulnerability to insult. Importantly, we use the outbred, wild-type CD1 mouse to generate our cultures, in order to examine the effect of the cytokine signature of AD progression in the absence of AD pathology, aiming to isolate the independent effect of the immune cues and eliminate any confounding factors caused by the overexpression of amyloid-related proteins in the 5xFAD mouse model.^75^

We stimulated cultures with recombinant IFN-γ, IP-10, and IL-9, in concentrations proportional to those measured in the 5xFAD hippocampus at 180 days of age and analyzed the expression of metabolism-associated genes (Methods). Metabolic and immune dysregulation are both early characteristics of the pre-clinical AD brain and are closely linked, contributing to an aggressive “immunometabolic” positive feedback loop that is capable of driving pathology progression.^87^ Cellular metabolism also provides a critical readout of cellular health, allowing fine-tuned assessment of the large “grey area” between live, healthy cells and cell death.

As Alzheimer’s disease is a neurodegenerative disease that selectively affects neurons, we first investigated the direct effect of the cytokine signature of AD progression on healthy primary neurons. We identified significantly differentially-expressed genes in neurons treated with the AD progression cytokine signature compared to vehicle-treated neurons (Supplementary Table 1) and analyzed their downstream protein interactions using the STRING database (Methods). Out of the 155 significantly differentially expressed genes in neurons following cytokine treatment, 80 were included in the main neuron STRING interaction network. The resulting neuron interaction network (Fig. 3A) demonstrated a wide variety of affected genes. Unsurprisingly, the genes most substantially effected by cytokine stimulation were associated with immune response. Genes involved in immune response were closely linked to genes involved in stress response, autophagy, and protein degradation processes, all of which are key dysregulated systems in AD.^21^ The network of affected neuronal transcripts also includes a down-regulated cluster of genes traditionally associated with the cell cycle. While neurons are post-mitotic cells, many of these genes have known functions in post-mitotic cells, including neurons: Plk1, cdc20, and kif2c are involved in the regulation of synapses, dendrites, and the cytoskeleton.^19,44,106^ Cdk9 and npm1 are implicated in cell survival,^51,63,67,69^ and ccna2 is associated with DNA repair in aging, rRNA homeostasis, and cellular senescence. ^3,36,100^ The down-regulation of this cluster of genes with treatment suggests increased neuronal and synaptic vulnerability to injury. Strikingly, the neuron network includes a grouping of primarily down-regulated genes involved in mitochondrial respiration, including subunits of every complex involved in the electron transport chain. The significantly down-regulated genes include the subunits Ndufa1 (complex I), Sdhc (complex II), Uqcr11 (complex III), Ndufa4 (Complex IV), and Atp5d (ATP synthase or complex V).

**Fig 3.**
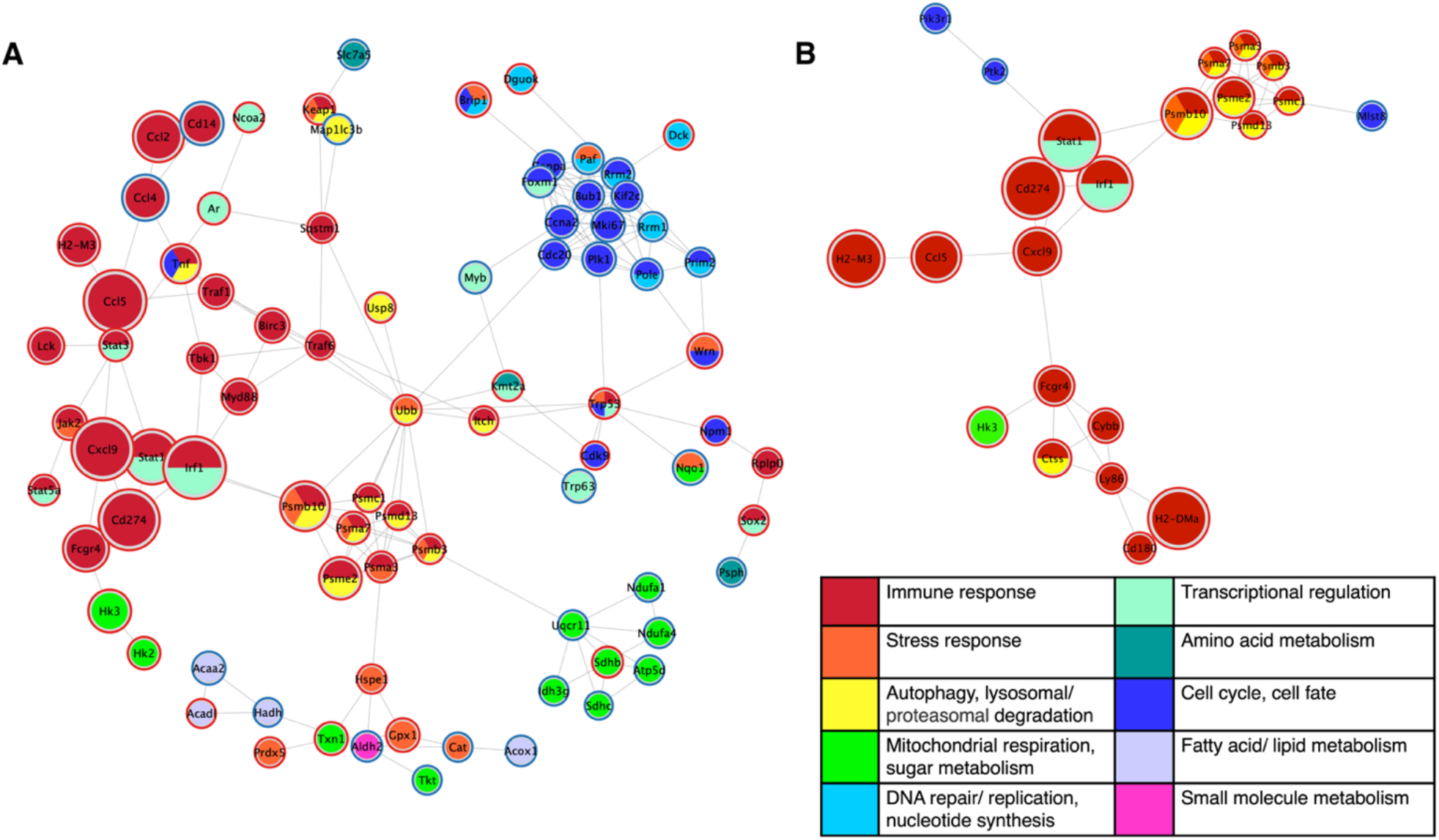
Cytokine signature of AD progression generates widespread gene expression changes in healthy neurons. STRING protein-protein interaction network of significantly differentially expressed genes in (A) neurons and (B) astrocytes in cytokine signature-treated primary cultures compared to vehicle controls. Individual genes, represented as nodes, are colored by their primary function(s) and sized according to the log2 fold change between treatments. Nodes are outlined in red or blue to indicate up- or down-regulation, respectively, with treatment. Network edges denote the STRING interaction score, with a shorter edge length indicating a stronger interaction. Thus, nodes that are closer together are more strongly linked.

The interaction network of differentially expressed genes (Supplementary Table 2) in cytokine-treated astrocytes compared to vehicle-treated astrocytes (Fig. 3B) demonstrated that fewer astrocyte genes were affected by the cytokine treatment than in neurons, and those that were affected are primarily involved in immune response. Out of the 54 significantly differentially expressed genes in astroctes following cytokine treatment, 23 were included in the main STRING interaction network. However, hk3, involved in glycolysis, was up-regulated in the astrocytes, which may indicate activation of a response to increase neuronal metabolic support.^2,65^ The astrocyte network also features a cluster of up-regulated proteasome related genes, many of which were also featured in the neuron network. Proteasomes are involved in the degradation of Aβ, and their dysfunction has been linked to the abnormal accumulation of pathological proteins in AD.^41,42,47^ The differential gene regulation indicates that AD-associated cytokine cues may activate astrocytes to bolster neuronal support functions, but the response to the AD signature was less robust than in neurons.

### Alzheimer’s disease-specific cytokine signature impairs neuronal metabolism

Treatment with the cytokine signature of AD progression resulted in marked down-regulation of sub-units in every electron transport chain complex. The resulting reduction of components responsible for mitochondrial respiration would result in significant impairment of neuronal oxidative phosphorylation. We stimulated primary neurons with the AD progression signature of IFN-γ, IP-10, and IL-9 and quantified parameters of mitochondrial function through oxygen consumption measurements on the Seahorse Extracellular Flux Analyzer (Agilent) using the Mito Stress Test Assay (Methods).

We found an overall decrease in oxygen consumption as a result of cytokine treatment compared to the vehicle control (Fig. 4). Basal respiration was significantly decreased in cytokine treated neurons (Fig. 4B, p-value: 0.031). Maximal respiration, induced by the mitochondrial membrane uncoupling agent FCCP, was similarly decreased with cytokine treatment (Fig. 4C, p-value= 0.012). ATP production also decreased following cytokine treatment, although this finding was not statistically significant (Fig. 4D, p-value: 0.065). Finally, the proton leak, the mitochondrial oxygen consumption not due to the action of ATP synthase, was also significantly reduced with treatment (Fig. 4E, p-value= 0.00021). Overall, the observed functional decrease in neuron mitochondrial respiration is in agreement with the gene expression findings of down-regulated electron transport chain complexes. A variety of subunits from all 5 complexes were down-regulated in the neuron cultures with IFN-γ, IP-10, and IL-9 treatment, and likely contributed the overall reduction in mitochondrial election transport chain performance that negatively impacts neuronal health and resilience.

**Fig 4.**
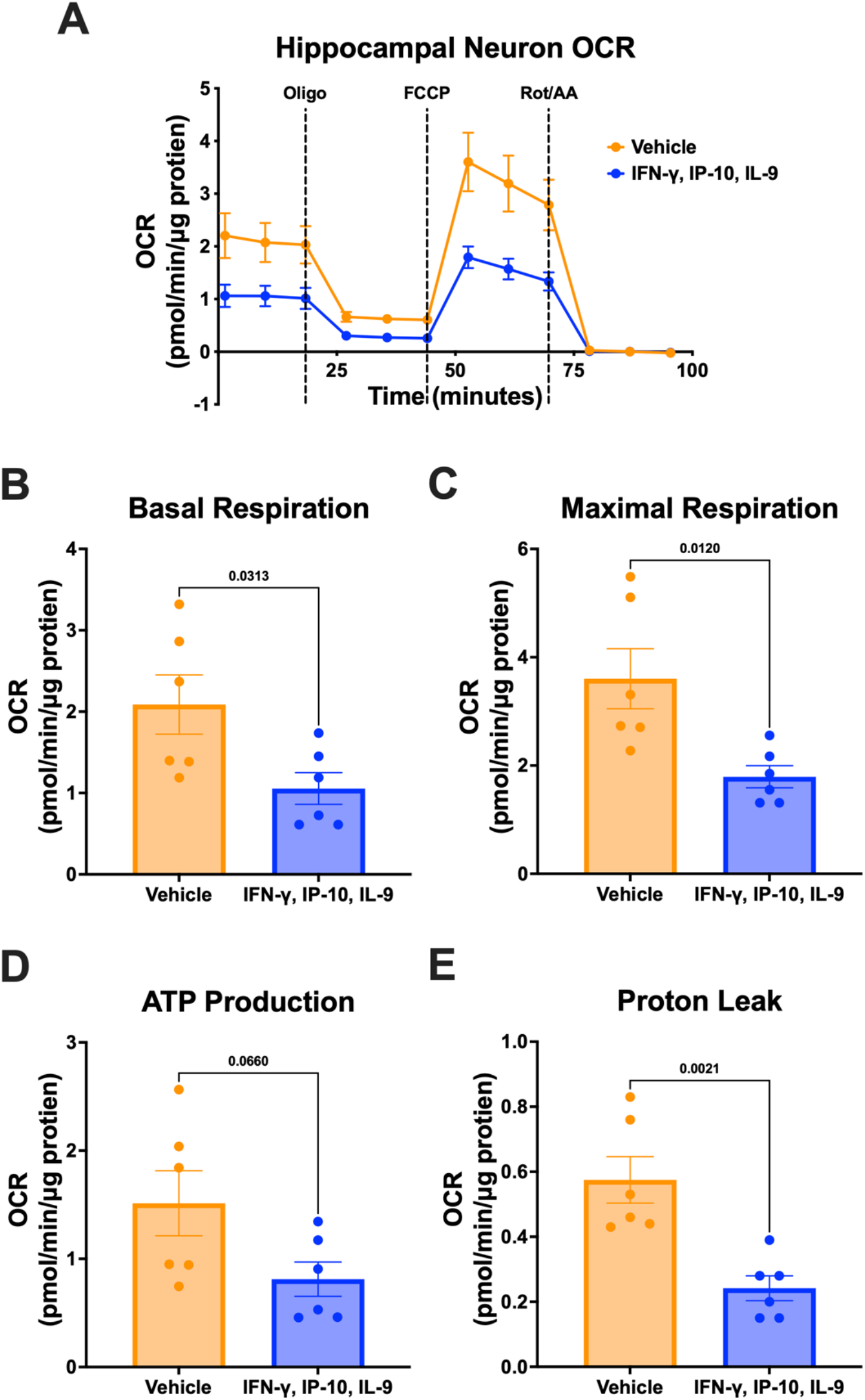
Wild-type primary hippocampal neurons treated with cytokine signature of AD progression exhibit decreased mitochondrial respiration. (A) Oxygen consumption during Seahorse Mito Stress Test following 72-hour vehicle (blue) or combination IFN-y, IP-10, IL-9 treatment (orange) and corresponding measurements of (B) basal respiration (p-value= 0.031), (C) maximal respiration (p-value= 0.012), (D) ATP production (p-value= 0.065), and (E) proton leak (p-value= 0.00021). P-values calculated by two-tailed student’s t-tests. Results generated from the average of 6 independent biological experiments with 5-10 technical replicates for each of treatment and control. Seahorse oxygen consumption rate measurements are displayed as mean ± standard error of the mean.

## Discussion

An incomplete understanding of AD etiology has prevented the development of successful strategies for prevention and treatment. Immune and metabolic dysfunction are among the earliest detectable pathologies in the disease, coinciding with or by some accounts preceding protein deposition.^40,81^ A greater understanding of the mechanisms underlying immune and metabolic dysfunction in AD has the potential to facilitate early interference to ameliorate disease. However, the complex and all-encompassing nature of entangled beneficial and harmful immune responses with intercellular communication and neuronal support functions complicate the identification of robust and effective targets. Recent systems biology approaches address this issue by characterizing the molecular-, cellular-, and tissue-level processes disrupted in AD, but have rarely gone beyond descriptive findings of altered cellular pathways.^21^ Here, we identified an AD-specific cytokine signature predictive of disease progression that defines the evolving immune state in the hippocampus, a primary site of neurodegeneration in AD and one of the earliest brain regions affected by the disease. Our signature was not exclusive to or built using a single cell type, instead representing the overall immunological milieu promoting AD progression. Notably, our signature does not include traditional pro-inflammatory cytokines associated with AD, such as TNF-α, IL-6, and IL-1β.^40^ While these cytokines are likely present in the tissue, their levels were below our threshold for inclusion in the model. While the individual members of the strong up-regulated signature we identified, IFN-γ, IP-10, and IL-9, have been linked to AD previously,^5,8,31,54,92^ with the possible exception of IP-10 they have not been appreciated as playing a strong role in AD progression. Further, the pathological role of these cytokines in neuronal metabolic dysfunction has never before been identified. We attribute our identifications of these cytokines to the multivariate nature of our model: the synergistic action of these cytokine cues drove the predictive power of our model, emphasizing the importance of considering emergent manifestation of network activities when identifying potential critical drivers of disease with possible strategic value as therapeutic targets.

Astrocytes and neurons are metabolically coupled, and dysfunction in astrocytes could predispose neurons to injury and death. We studied the independent effects of our cytokine signature of AD progression on both cell types to identify potential pathways of neuronal injury and subsequent contribution to disease. Receptors for IFN-γ, IP-10, and IL-9 are found on both neurons^27,53,85,97^ and astrocytes,^11,26,37,107^ supporting the study of their combined effect on each of these cell types. While AD signature-stimulated changes in astrocyte gene expression were limited and primarily related to immune response, we identified numerous differentially expressed genes in neurons with known AD implications, such as regulation of synapse plasticity and dendrite projection and tryptophan and mitochondrial metabolism. We note that these assays were specifically performed in healthy, wild-type primary neuron cultures in the absence of any pre-existing AD pathology. Our objective was to determine the capacity of the AD progression cytokine signature to affect neuron health and viability, which is most cleanly demonstrated in the absence of any other AD-promoting factors. The fact that treatment with the cytokine signature altered expression of genes previously linked to Alzheimer’s disease, in the direction that would promote disease and absent of any other AD-related factors present, demonstrates the power of our model to extract patterns of effectors with discernible value for mechanistic insight. These insights have broad potential toward predicting diverse types of outcomes in AD and AD risk; we have previously used the partial least squares method to identify brain cytokine signatures predictive of systemic metabolism^25^ and to discover a previously unknown neurotoxic effect of VEGF in the presence of Aβ aggregates.^96^

The suppression of neuronal mitochondrial metabolism by the cytokine signature of AD progression indicates that, at baseline under normal energy demands, neuronal metabolism in this immune environment does not function efficiently and would not be able to adequately respond to an increased energy demand when presented with a disease insult such as AD-related proteinopathy. The energy demands of the brain are great, far exceeding other organs of the body, and are primarily achieved through glucose metabolism.^18^ Impaired metabolic capacity in neurons could predispose to neuronal injury and neurodegeneration by an inability to sustain the energetic demand of synapse activity and other neuronal functions while under stress.^18^

The cytokines comprising our signature of AD progression have not previously been connected to a impaired metabolism. However, dysregulated immune signaling and metabolic impairment are both well-known features of AD etiology, and their synergistic connection has been established in other tissues and diseases such as cancer and type 2 diabetes.^49,61^ In AD, glial activation and the accompanying cytokine release causes downstream metabolic stress,^20^ which is largely characterized by an increase in reactive oxygen species (ROS) and decreased ETC activity.^64,71,77,81^ While not directly connected to neuroinflammation or cytokine signaling, multiple subunits of mitochondrial ETC complexes are down-regulated in the AD brain,^87^ including in the hippocampus,^76^ temporal cortex,^58^ and frontal cortex.^1,50^ Outside of the brain, cytokines have been implicated in the reduced expression of ETC complex genes and impaired mitochondrial respiration in cultures of other cell-types, such as HepG2 cells (a cell line originating from a hepatocellular carcinoma),^43,90^ cardiomyocytes,^103^ and hepatocytes^9,46^.

## Conclusion

Here, we identified a hippocampal cytokine signature of AD progression that is predictive of advancing disease in the 5xFAD mouse model of AD. Exposure to this signature of AD progression was sufficient to initiate mitochondrial dysfunction in primary neuron cultures that was similar to that previously observed in AD, but in the complete absence of AD pathology. The resulting decrease in mitochondrial respiration predisposes neurons to injury and in the presence of a stressor such as proteinopathy could lead to energetic insufficiency and, ultimately, cell death. The cytokines comprising our signature of AD progression have not previously been connected to mitochondrial dysfunction, highlighting the power of a systems biology approach that considers the inter-relatedness of immune signaling. Overall, our findings support the existence of an immunometabolic feedback loop driving neuronal vulnerability in AD. Further investigation is needed to evaluate the relevance of the signature in human AD rather than the mouse model used here, particularly in the presence of tau pathology, which is absent in the 5xFAD model. However, the interdependency and redundancy of cytokine signaling, as well as the differences between mice and humans, make it likely that there is a corresponding human signature capable of driving mitochondrial dysfunction.

## Acknowledgements

This work was supported by R21AG068532 from the National Institute on Aging (EAP) and start-up funds from Penn State College of Medicine Departments of Neurosurgery and Pharmacology (EAP). MKK and DCC are supported by training fellowship T32NS115667 from the National Neurological Disorders and Stroke. RMF is supported by predoctoral fellowship F31AG071131 from the National Institute on Aging. We would like to thank the Penn State Genomics Core Facility (University Park, PA) for facilitating the use of the NanoString platform and the NanoString Proof of Principle Lab for their guidance on data processing.

## Abbreviations

AD: Alzheimer’s disease
Aβ: Amyloid beta
ETC: Electron Transport Chain
NRF: Nuclear Respiratory factor
OCR: Oxygen Consumption Rate
PLS-DA: Partial least squares discriminant analysis
PLSR: Partial least squares regression
ROS: Reactive Oxygen Species
VIP: Variable importance in projection

---

**Supplementary Table 1:**
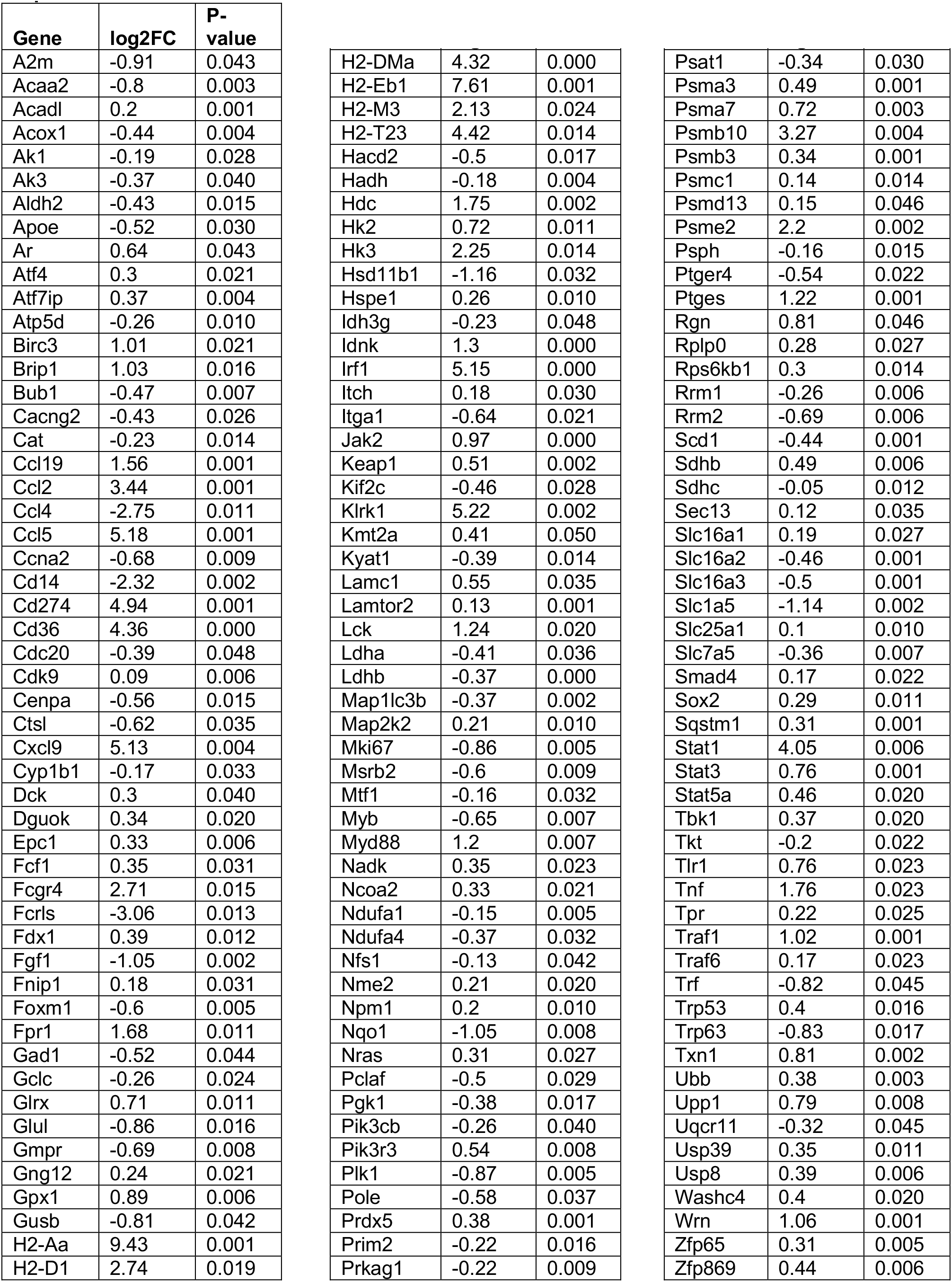
Significantly differentially expressed genes of cytokine-treated neurons compared to vehicle-treated neurons.

**Supplementary Table 2:** Significantly differentially expressed genes of cytokine-treated astrocytes compared to vehicle-treated astrocytes.

| Gene | log2FC | P-value |
| --- | --- | --- |
| Ada | 0.73 | 0.049 |
| Akt3 | 0.48 | 0.002 |
| Cab39 | -0.22 | 0.025 |
| Ccl2 | 2.88 | 0.037 |
| Ccl5 | 3.55 | 0.016 |
| Cd180 | 0.7 | 0.020 |
| Cd274 | 5.36 | 0.000 |
| Ctss | 1.63 | 0.003 |
| Cxcl9 | 3.21 | 0.017 |
| Cybb | 1.2 | 0.002 |
| Dnajc14 | -0.15 | 0.043 |
| Fcgr4 | 2.02 | 0.001 |
| Fgf1 | -0.85 | 0.003 |
| Gns | 0.23 | 0.040 |
| Gsk3b | -0.29 | 0.020 |
| H2-Aa | 8.65 | 0.001 |
| H2-D1 | 5.79 | 0.000 |
| H2-DMa | 5.32 | 0.000 |
| H2-Eb1 | 6.87 | 0.001 |
| H2-M3 | 4.61 | 0.000 |
| H2-T23 | 4 | 0.003 |
| Hexa | 0.27 | 0.003 |
| Hk3 | 2.07 | 0.008 |
| Hspa2 | 0.52 | 0.011 |
| Hspa4 | -0.39 | 0.001 |
| Idh2 | -0.49 | 0.036 |
| Idnk | 1.31 | 0.007 |
| Irf1 | 4.39 | 0.001 |
| Itgb2 | 0.91 | 0.041 |
| Ldha | -0.46 | 0.025 |
| Ly86 | 0.54 | 0.045 |
| Map2k1 | 0.1 | 0.028 |
| Mlst8 | -0.39 | 0.026 |
| Mycn | -0.5 | 0.021 |
| Ndufa3 | 0.17 | 0.028 |
| Pik3r1 | -0.4 | 0.035 |
| Prkcg | -0.63 | 0.032 |
| Psma3 | 0.46 | 0.008 |
| Psma7 | 0.55 | 0.009 |
| Psmb10 | 3.73 | 0.000 |
| Psmb3 | 0.35 | 0.006 |
| Psmc1 | 0.17 | 0.013 |
| Psmd13 | 0.16 | 0.042 |
| Psme2 | 2.17 | 0.000 |
| Ptk2 | -0.25 | 0.015 |
| Pycr1 | 0.59 | 0.021 |
| Rbks | -0.34 | 0.025 |
| Sdha | -0.25 | 0.050 |
| Slc25a1 | 0.23 | 0.014 |
| Sqstm1 | 0.29 | 0.044 |
| Stat1 | 5.3 | 0.000 |
| Tyms | -0.44 | 0.033 |
| Washc4 | 0.3 | 0.000 |
| Wrn | 0.53 | 0.048 |
| Xdh | 1.67 | 0.006 |
| Zfp457 | -0.67 | 0.001 |

